# Dynamic meta-analysis: a method of using global evidence for local decision making

**DOI:** 10.1101/2020.05.18.078840

**Authors:** Gorm E. Shackelford, Philip A. Martin, Amelia S. C. Hood, Alec P. Christie, Elena Kulinskaya, William J. Sutherland

## Abstract

Meta-analysis is often used to make generalizations across all available evidence at the global scale. But how can these global generalizations be used for evidence-based decision making at the local scale, if only the local evidence is perceived to be relevant to a local decision? We show how an interactive method of meta-analysis — dynamic meta-analysis — can be used to assess the local relevance of global evidence. We developed Metadataset (www.metadataset.com) as an example of dynamic meta-analysis. Using Metadataset, we show how evidence can be filtered and weighted, and results can be recalculated, using dynamic methods of subgroup analysis, meta-regression, and recalibration. With an example from agroecology, we show how dynamic meta-analysis could lead to different conclusions for different subsets of the global evidence. Dynamic meta-analysis could also lead to a rebalancing of power and responsibility in evidence synthesis, since evidence users would be able to make decisions that are typically made by systematic reviewers — decisions about which studies to include (e.g., critical appraisal) and how to handle missing or poorly reported data (e.g., sensitivity analysis). We suggest that dynamic meta-analysis could be scaled up and used for subject-wide evidence synthesis in several scientific disciplines (e.g., agroecology and conservation biology). However, the metadata that are used to filter and weight the evidence would need to be standardized within disciplines.

## Introduction

Meta-analysis is often used to make generalizations about interventions, such as agricultural practices or medical treatments (Gurevitch *et al.* 2018). It can be difficult to make generalizations if interventions have different effects in different contexts. For example, a meta-analysis of conservation agriculture found beneficial effects in hotter, drier climates, but not in colder, wetter climates (Steward *et al.* 2018). Therefore, it can be difficult to use meta-analysis to make decisions about interventions in a specific context, unless the results are known to be generalizable to that specific context.

What is needed is a method of meta-analysis that enables decision makers to answer the question, “How effective is this intervention in my specific context?” (Wang, Moss & Hiller 2005; Burford *et al.* 2013; Avellar *et al.* 2016). Subgroup analysis and meta-regression (Borenstein *et al.* 2009) are standard methods of meta-analysis that can be used to answer this question, but only if the researchers who produce the meta-analysis ask the same question as the decision makers who use a meta-analysis. In the above example of conservation agriculture (Steward *et al.* 2018), researchers used meta-regression to ask, “How effective is conservation agriculture in different climates?” But decision makers may want to ask, “How effective is conservation agriculture in my climate or in my country?” Researchers may not publish an answer to this specific question, not only because they do not know which variables will define the context for different decision makers, but also because they do not have the time and space to analyse and publish the results for all combinations and permutations of context-defining variables. Instead, researchers may only publish an answer to a more generic question.

The lack of context-specific evidence is a problem in evidence-based decision making (Christie *et al.* in press; Innvær *et al.* 2002; Cook, Possingham & Fuller 2013). One solution to this problem is to commission new research and/or new reviews that exactly match the local context (e.g., “co-production” of knowledge), but that takes time and money and may be impractical or impossible for many decisions. Another solution is to assess the relevance of existing research that does not exactly match the local context (e.g., “co-assessment” of knowledge (Sutherland, Shackelford & Rose 2017)). Relevance includes both “applicability” and “transferability” (Wang, Moss & Hiller 2005). *Transferability* is the extent to which an intervention would have the same effect in a different context (e.g., conservation agriculture might have a different effect in a different climate). *Applicability* is the extent to which an intervention would be feasible in a different context (e.g., conservation agriculture might not be feasible in an area without access to herbicides or seed drills). We use these terms as defined above (in the sense of Wang, Moss & Hiller 2005), but we note that applicability, transferability, external validity, and generalizability are sometimes used interchangeably and are sometimes used in somewhat different senses (Burchett, Umoquit & Dobrow 2011; Burford *et al.* 2013). Here, we focus on transferability, but we also discuss applicability.

It has been suggested that “research cannot provide an exact match to every practitioner’s circumstances, or perhaps any practitioner’s circumstances because environments are dynamic and often changing, whereas completed research is static” (Avellar *et al.* 2016). A partial solution to this problem could be to make research more dynamic, by enabling decision makers to interact with it. For example, decision makers could filter a database of research publications, to find studies that are more relevant to their circumstances, or they could weight these studies by relevance to their circumstances. Several methods of interactive evidence synthesis have already been developed. For example, interactive evidence maps enable users to filter research publications by country (e.g., McKinnon *et al.* 2015). Decision-support systems enable users to weight evidence by value to stakeholders (e.g., Shackelford *et al.* 2019). However, as far as we are aware, there are no tools that enable users to filter and weight the studies in a meta-analysis, and thereby to answer the question, “How effective is this intervention in my specific context?” Therefore, we developed a tool for this purpose, and here we show how this tool could be used to assess the local relevance of a global meta-analysis in agroecology.

This tool is an example of a method that we refer to as *dynamic meta-analysis*. This term has been used in different disciplines and in different senses (cf. Garamszegi, Nunn & McCabe 2012; Maki, Cohen & Vandenbergh 2018; Becker *et al.* 2020), and sometimes in the sense of a *living systematic review* that can be dynamically updated by researchers (Elliott *et al.* 2014; Bergmann *et al.* 2018), instead of a meta-analysis that can be dynamically filtered and weighted by users. However, as far as we are aware, dynamic meta-analysis has not been defined as a method, and we define it here.

## Methods

### Dynamic meta-analysis

As we define it here, dynamic meta-analysis is a method of interactively filtering and weighting the data in a meta-analysis. The diagnostic feature of a dynamic meta-analysis is that it takes place in a *dynamic* environment (e.g., a web application), not a *static* environment (e.g., a print publication), and this enables users to interact with it. Dynamic meta-analysis includes *subgroup analysis* and/or *meta-regression* (Borenstein *et al.* 2009). These are standard methods in meta-analysis, and they are used to calculate the results for a subset of the data, either by analysing only that subset (subgroup analysis) or else by analysing all of the data but calculating different results for different subsets, while accounting for the effects of other variables (meta-regression). The variables that define these subsets can include country, climate type, soil type, study design, or any other metadata that can be used to define relevance. In a dynamic meta-analysis, users filter the data to define a subset that is relevant to them, and then the results for that subset are calculated, using subgroup analysis and/or meta-regression.

Dynamic meta-analysis also includes *recalibration* (Kneale *et al.* 2019), which is a method of weighting studies based on their relevance. With recalibration, users can consider a wider range of evidence — not only the data that is completely relevant, but also the data that is partially relevant. Recalibration may be the only option, if no evidence exists that is completely relevant.

Dynamic meta-analysis also includes elements of *critical appraisal* (i.e. deciding which studies should be included in the meta-analysis, based on study quality) and *sensitivity analysis* (i.e. permuting the assumptions of a meta-analysis, to test the robustness of the results). Critical appraisal and sensitivity analysis are typically performed by systematic reviewers (e.g., see the Collaboration for Environmental Evidence (CEE) 2018 for standard methods), but dynamic meta-analysis enables decision makers to participate in both critical appraisal and sensitivity analysis.

For example, decision makers may want to include or exclude a controversial study. Or they may want to include studies that are relevant to their local context, even though these studies are lower-quality, if higher-quality studies are not available (McGill *et al.* 2015). For example, if decision makers are looking for conservation studies on a specific biome or taxon, higher-quality studies may not be available (Christie *et al.* in press). In some forms of evidence synthesis, lower-quality studies are excluded from the evidence base before they can be considered by decision makers (e.g., *best evidence synthesis* (Slavin 1986)), but in a dynamic meta-analysis these studies can be included in the evidence base and tagged with metadata, so that decision makers can consider these studies for themselves.

It may also be important to include all studies, regardless of study quality, if study quality is related to study results. For example, in a review of forest conservation strategies, lower-quality studies were more likely to report negative results (Burivalova *et al.* 2019). By comparing the results of different analyses that are based on different studies or different assumptions (e.g., different methods for handling missing data), users can test the sensitivity of the results to these different assumptions (sensitivity analysis).

### Metadataset: a website for dynamic meta-analysis

We developed Metadataset (www.metadataset.com) to show how dynamic meta-analysis can be used. Metadataset is a website that provides two methods of interactive evidence synthesis: (1) browsing publications by intervention, outcome, or country (using interactive evidence maps) (Figure 1), and (2) filtering and weighting the evidence in a dynamic meta-analysis (Figure 2). Supplementary File 1 is a video that shows how Metadataset can be used.

**Figure 1.**
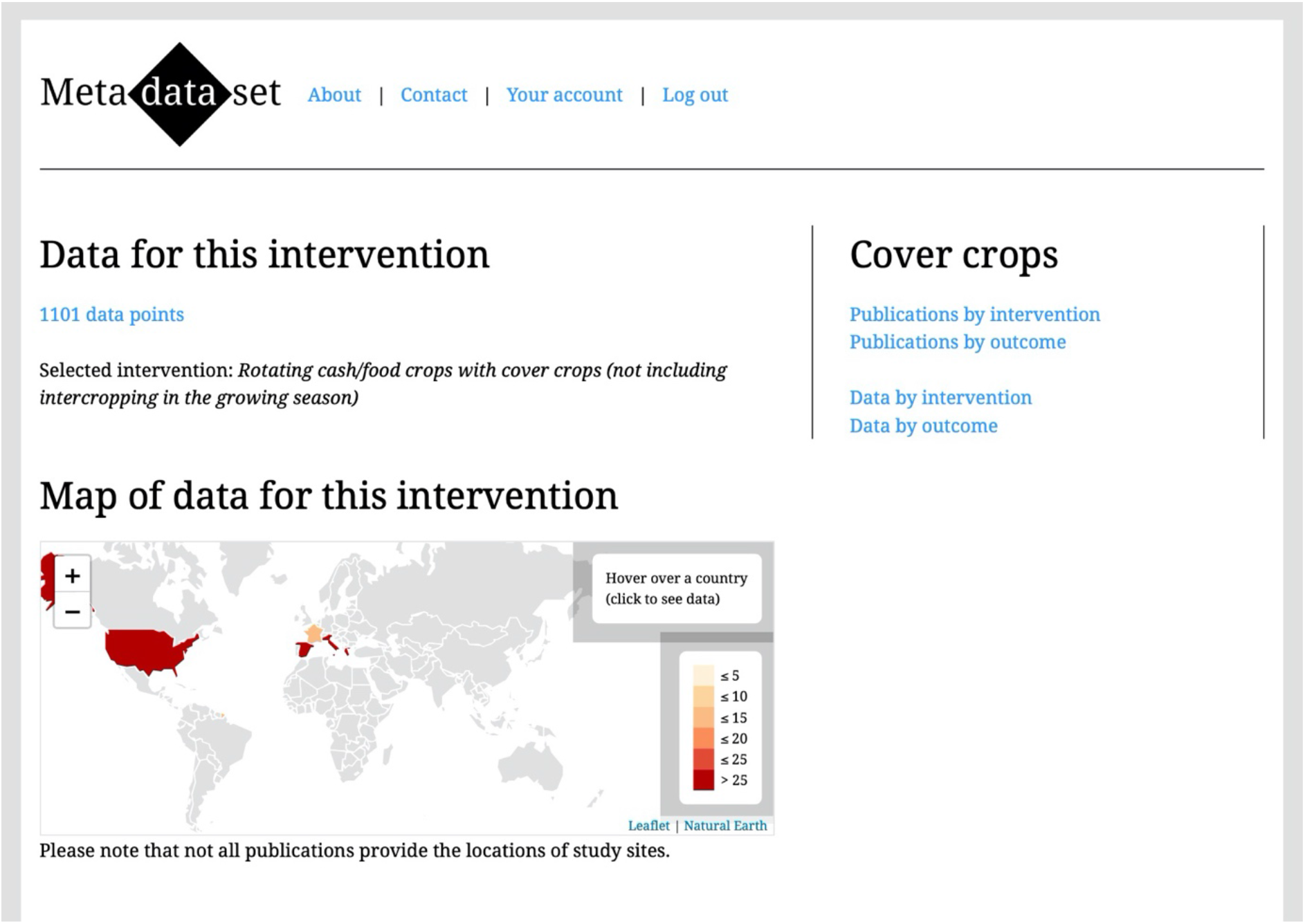
A screenshot from Metadataset (www.metadataset.com) that shows an interactive evidence map.

**Figure 2.**
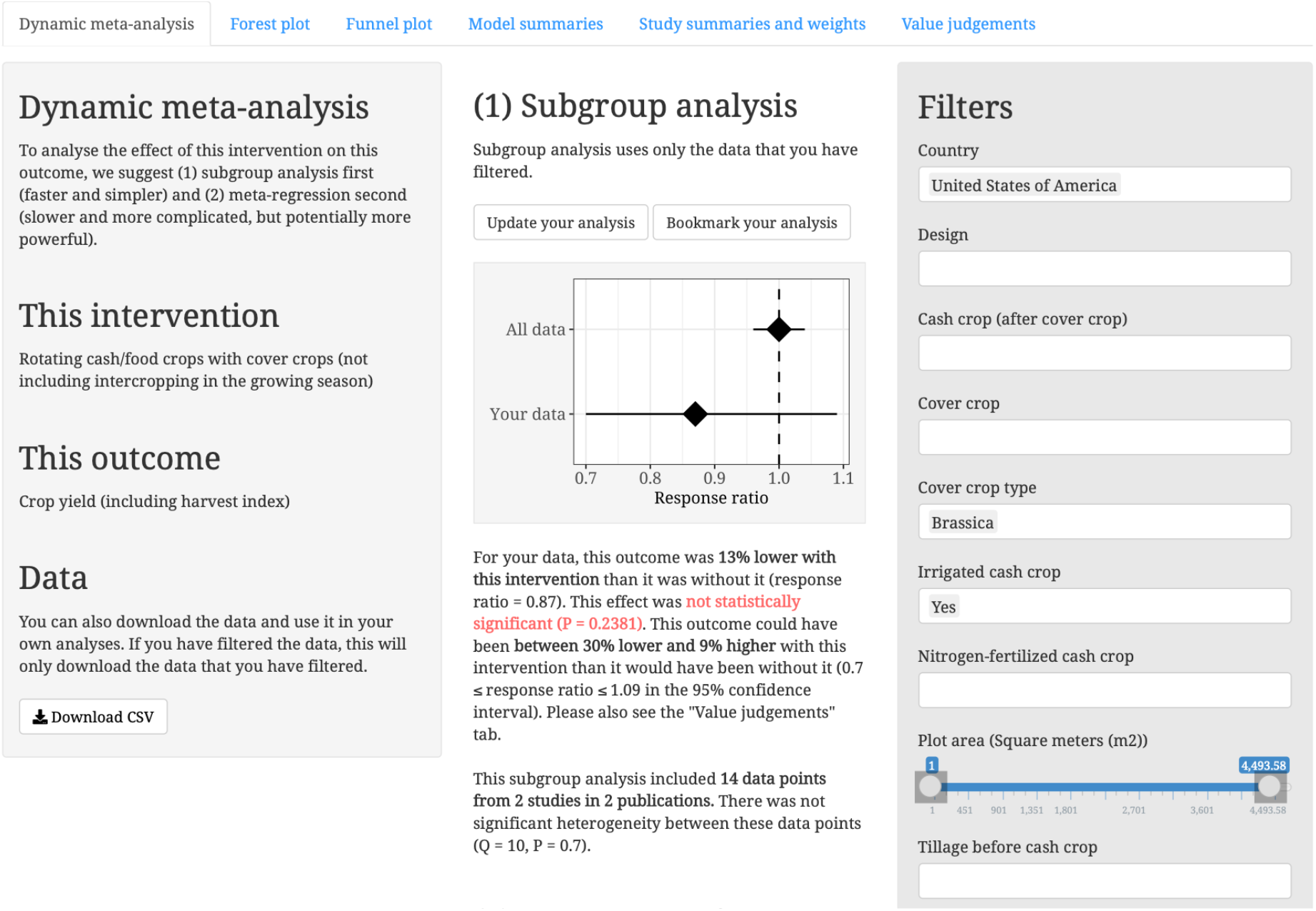
A screenshot from Metadataset (www.metadataset.com) that shows a dynamic meta-analysis.

At present, Metadataset has evidence on two subject areas: (1) agriculture, which includes data from a meta-analysis of cover crops in Mediterranean climates (Shackelford, Kelsey & Dicks 2019) and a systematic map of cassava farming practices that is a work in progress (Shackelford *et al.* 2018); and (2) invasive species, which includes a systematic review of management practices for invasive plants that is also a work in progress (Martin *et al.* 2020). However, we plan to expand Metadataset to other subject areas, and we welcome collaborations. Here we focus on cover crops in Mediterranean climates as an example of dynamic meta-analysis.

Cover crops are often grown over the winter, as an alternative to bare soil or fallow, and cash crops are grown over the following summer. Shackelford et al. (2019) analysed the effects of cover crops on ten outcomes (e.g., cash crop yield and soil organic matter) and recorded the metadata that we use here for subgroup analysis and meta-regression (e.g., country, cover crop type, fertilizer usage, and tillage). Shackelford et al. (2019) presented some subgroup analyses (e.g., legumes vs non-legumes as cover crop types), but noted the problem of not being able to report all combinations of subgroups that might be of interest to a reader (e.g., legumes in California, without synthetic fertilizer). We entered their data into Metadataset, to show how dynamic meta-analysis is a solution to this problem.

Metadataset uses hierarchical classifications of interventions and outcomes, so that the evidence can be analysed at different levels of resolution. For example, users can analyse the effects of one intervention (e.g., growing cover crops) on one outcome (e.g., soil organic matter) or multiple outcomes (e.g., all soil outcomes, including soil organic matter, but also soil erosion, soil nutrients, etc.). Not only can users decide which studies should be included in the meta-analysis, but they can also decide which interventions and outcomes should be included. Please see the Metadataset website for examples of these classification systems (e.g., https://www.metadataset.com/subject/cassava/browse-by-intervention/publications/).

The Metadataset website is built on two separate web frameworks: (1) the Django framework for Python (www.djangoproject.com), and (2) the Shiny framework for R (https://shiny.rstudio.com). Using the Django app, researchers can screen publications for inclusion in evidence maps and can tag these publications with interventions, outcomes, and other metadata. They can then enter the data that will be used for dynamic meta-analysis (e.g., the mean values for treatment groups and control groups, standard deviations, numbers of replicates, and *P*-values), and they can write paragraphs that summarize each study.

Users can browse this evidence by intervention, outcome, or country, to find relevant publications and/or datasets. They can then click a link to the Shiny app, to interact with their selected datasets using dynamic meta-analysis. The code is open source (Django app: https://github.com/gormshackelford/metadataset), Shiny app: https://github.com/gormshackelford/metadataset-shiny), and the data is open access (the data can be downloaded in CSV files via the Shiny app). Metadataset was developed as part of Conservation Evidence (www.conservationevidence.com) and BioRISC (the Biosecurity Research Initiative at St Catharine’s College, Cambridge; www.biorisc.com).

### Methods for dynamic meta-analysis on Metadataset

The Shiny app uses the methods from Shackelford et al. (2019) to calculate the mean effect size of an intervention as the *log response ratio*. The *response ratio* is the numerical value of an outcome, measured with the intervention, divided by the numerical value of an outcome, measured without the intervention. The natural logarithm of the response ratio (the log response ratio) is typically used for meta-analysis (Hedges, Gurevitch & Curtis 1999). Using the rma.mv function from the metafor package in R (Viechtbauer 2010), the Shiny app fits a mixed-effects meta-analysis that accounts for non-independence of data points (for example, multiple data points within one study, within one publication) by using random effects (e.g., “random ~ 1 | publication/study” in the rma.mv function in metafor). Users can select, deselect, and/or adjust settings for missing or poorly reported data. For example, there are settings for imputing the variance of studies with missing variances (using the mean variance), approximating the variance of studies with missing variances (based on their *P*-values; see Shackelford *et al.* (2019)), and excluding outliers. Please see Supplementary File 2 for more information on methods and settings.

Users can filter the data (e.g., they can select “Brassica” from the filter for “Cover crop type”), and then they can use subgroup analysis and/or meta-regression to recalculate the results (see Figure 2). They can view forest plots and funnel plots of their filtered data and read the paragraphs that summarize the studies that are included in their analyses. They can also assign a weight to each study, based on its relevance to their decision-making context. It has been suggested that a ratio of 5:4 (one “deciban”) is the smallest difference in the weight of evidence that is perceptible to humans (Good 1985). Therefore, we allow users to assign weights on a scale from 0 to 1, in increments of 0.1, without allowing weights that are overly precise and beyond human perception (e.g., a ratio of 1:0.99).

The standard method in meta-analysis is to weight each study by the inverse of its variance, so that studies with smaller variances have larger weights. To weight each study not only by the inverse of its variance, but also by its relevance (assigned by the user), we specify a weight matrix, W, using the following equation:

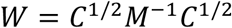

*C* is a diagonal matrix of relevance weights (one weight for each study, assigned by the user, with a default weight of 1), and *M* is the default variance-covariance matrix in metafor (please see Supplementary File 3 for an example). The default weight matrix in metafor is the inverse of *M*, and here we multiply it by the square-root of *C*, our relevance matrix, twice (effectively multiplying *M* by *C*, but maintaining a symmetrical weight matrix). With a relevance weight of 1 for each study (the default setting), this has no effect on the weight matrix, and thus it is also possible for users to fit a model with inverse-variance weights. However, with a relevance weight of less than 1, a study has less effect on the mean effect size. We use this method as an example of recalibration, in the sense of Kneale et al. (2019). Kneale et al. (2019) provided an example of weighting studies in a meta-analysis, based on the similarity of these studies to different decision contexts, but they noted their method was provisional. Our method of modifying the weight matrix is also provisional. However, we think it is useful as an example of recalibration. Similar methods for using study-quality weights have been implemented in other meta-analyses, but it has been suggested that these methods also need further research (Stone *et al.* 2020).

After selecting filters and doing a subgroup analysis, with or without recalibration, users can also do a meta-regression. The Shiny app fits a model in metafor, as before, but with all of the selected filters and all of their two-way interactions as moderators. For example, if the user selects a filter for “Country” and a filter for “Cover crop type” then we fit a metafor model with “mods = ~ Country + Cover.crop.type + Country:Cover.crop.type”. We then use the MuMIn package in R (Bartoń 2009) to fit all possible combination of these moderators (e.g., a model without the two-way interaction term, or a model without any moderators). We then use the “best” model (with the lowest AICc score) to get the model predictions for the filters that the user selected (e.g., the results for brassicas in the USA; please see Supplementary File 3 for an example). We show these results to the user, together with the results from the subgroup analysis for the same filters. If one or more of the filters were not included in the meta-regression model, then we show a warning.

If a dynamic meta-analysis is done at a high level in the hierarchy of outcomes (e.g., soil), then it may include multiple outcomes (e.g., soil organic matter, soil nitrate leaching, and soil water content), and therefore the user may need to decide whether it is better for the intervention to cause an increase or a decrease in each outcome. Without doing this, the overall effect size will not be meaningful across multiple outcomes. There are settings for this in the Shiny app (on the “Value judgements” tab). For example, the user could decide that an increase in soil organic matter and soil water content, but a decrease in soil nitrate leaching, would be good outcomes in their context. The user would then select “decrease is better” for soil nitrate leaching. The Shiny app would then invert the response ratio for that outcome, so that a positive effect size would represent a good outcome across all outcomes.

### An example of dynamic meta-analysis on Metadataset

To show how Metadataset can be used for dynamic meta-analysis, we imagine a scenario in which a hypothetical user searches for evidence on cover crops that are brassicas (e.g., mustard or rapeseed) on irrigated farms in California. Brassicas do not fertilize the soil as legumes do (by fixing nitrogen), and their negative effects on soil fertility (including allelochemicals that poison the soil for other plants) could have negative effects on the yields of the cash crops that are grown over the following summer, even if they do successfully suppress weeds over the winter. Thus, there is reason to believe that the evidence on cover crops in general may not be transferable to specific cover crops, such as brassicas or legumes, which have different effects on the soil (Shackelford, Kelsey & Dicks 2019). We show how this hypothetical user filters and weights the evidence on Metadataset (also see Table 1 for a summary of the steps in a dynamic meta-analysis, based on this scenario; see Supplementary File 1 for a video that shows this scenario; and see Supplementary File 3 for offline R code that reproduces the online results from the Shiny app).

**Table 1.** An example of the steps in a dynamic meta-analysis.

| Step | Action | Result |
| --- | --- | --- |
| **1. Meta-analysis** of all studies | Browse Metadataset by intervention and/or outcome, make selections, and click “Start your analysis” in the Shiny app | Crop yield: 0% different after cover crops (non-significant) |
| **2. Subgroup analysis** of selected studies | Select filters and then click “Update your analysis” | Crop yield: 13% lower after cover crops (non-significant) (brassicas in the USA, with irrigated cash crops) |
| **3. Meta-regression** of all studies, with results for selected studies | With the same selections, click “Meta-regression” | Crop yield: 9% lower after cover crops (significant) (brassicas in the USA, but irrigation was not included in the best model) |
| **4. Recalibration** of selected studies | Move the sliders on the tab for “Study summaries and weights” and then click “Update your analysis” | Crop yield: 17% lower (significant) (brassicas in the USA, with irrigated cash crops, and with a relevance weight of 0.5 assigned to one study) |
| **5. Sensitivity analysis** | Permute the settings (e.g., methods for handling missing data) and then compare the results | Crop yield: significantly lower than 0% |

The evidence on cover crops includes 57 publications from 5 countries: France (2 publications), Greece (2), Italy (24), Spain (9), and the United States of America (20) (https://www.metadataset.com/subject/cover-crops/). Browsing the data by outcome, this user finds the hierarchical classification of outcomes. She clicks “filter by intervention” for one outcome (“10.10.10. Crop yield”) and she sees that there are 316 data points for this outcome. She clicks an intervention (“Rotating cash/food crops opens. with cover crops”), and the Shiny app opens

To see the results for all 316 data points in the Shiny app, she deselects the option for “Exclude rows with exceptionally high variance (outliers)” and then she clicks “Start your analysis” to start a dynamic meta-analysis for her selected intervention and outcome (Step 1 in Table 1). Based on all 316 data points from 38 publications, cover crops do not have significant effects on cash crop yields (response ratio = 1; *P* = 0.9788; cash crop yields are 0% different with cover crops than they are without cover crops, with a 95% confidence interval from 4% lower to 4% higher).

However, these are the generic results for all of the global evidence. To find results that are transferable to her specific context, she filters the evidence (Step 2 in Table 1). She selects “United States of America” from the filter for “Country”, “Brassica” from the filter for “Cover crop type”, and “Yes” from the filter for “Irrigated cash crop”. She then clicks “Update your analysis” to see the subgroup analysis for these filters (Figure 2). Based on 14 data points from 2 publications (the only publications in which the cover crops were brassicas, grown in the USA, followed by irrigated cash crops), cash crop yields are lower after cover crops, but not significantly lower (13% lower, with a 95% confidence interval from 30% lower to 9% higher; *P* = 0.2381).

She clicks “Meta-regression” to see if the results from this subgroup analysis are relatively similar to the results from the meta-regression (Step 3 in Table 1). In the meta-regression, cash crop yields are significantly lower after cover crops (9% lower, with a 95% confidence interval from 12% lower to 5% lower; *P* < 0.0001). This is not surprising, since meta-regression is potentially more powerful statistically than subgroup analysis (it uses all of the data, and it potentially produces better estimates of variance). However, she sees a warning that one of her selected filters (“Irrigated cash crop”) did not have a significant effect on this outcome (i.e. this moderator was not included in the “best” meta-regression model, with the lowest AICc). She deselects this filter and clicks “Update your analysis”. There are now 30 data points from 3 publications in the subgroup analysis, and yields are now significantly lower (*P* = 0.0436). So far, it seems that the global evidence is not transferable to her local conditions (neutral effects vs negative effects on cash crop yields). However, she has found some evidence that seems transferable, and she has recalculated the results for this evidence, using subgroup analysis and meta-regression.

She clicks the tab for “Study summaries and weights” to see the paragraphs that summarize each of these three studies (Figure 3). She sees one study on maize, one on tomatoes, and one on beans. Tomatoes are less applicable in her interests (she is mostly interested in grains or pulses as cash crops), so she sets a relevance weight of 0.5 for the study on tomatoes. She then returns to the tab for “Dynamic meta-analysis” and clicks “Update your analysis” to see the effects of this recalibration (Step 4 in Table 1). The results are still negative, but slightly more significant (*P* = 0.0224).

**Figure 3.**
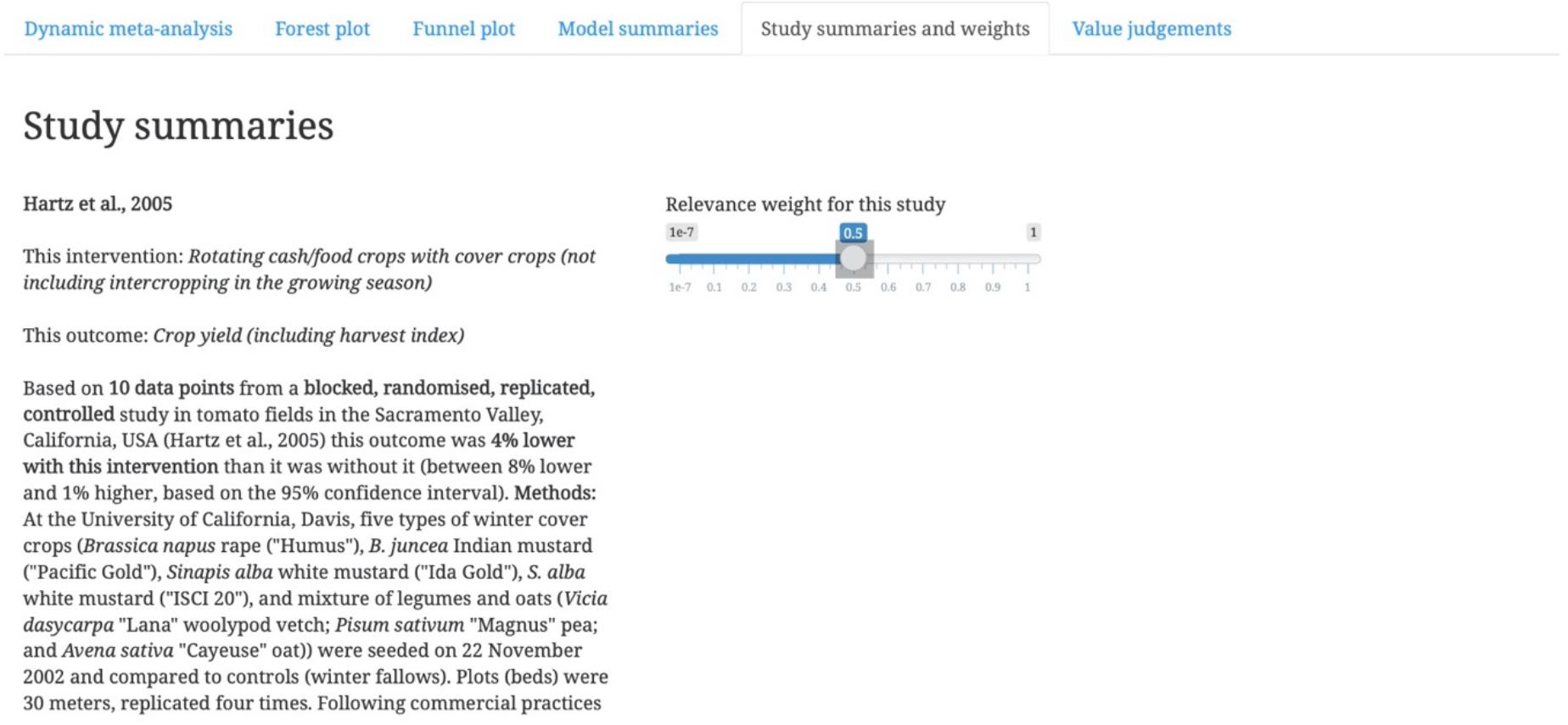
A screenshot from Metadataset (www.metadataset.com) that shows a method of recalibration in a dynamic meta-analysis. Users can adjust the weight of a study, based on its relevance to their context.

She then considers the sensitivity of these results by permuting the settings. For example, there are several options for handling missing data, and these can be selected, deselected, and/or adjusted for sensitivity analysis (Step 5 in Table 1). Deselecting the option for “approximate the variance of the log response ratio” (below the filters), the result is still significantly negative. Permuting several other options (e.g., the sliders for assumed *P*-values), this result seems to be robust (all of the results are significantly negative).

She reaches the conclusion that cover crops could have negative effects on cash crop yields in her local conditions (brassicas as cash crops on irrigated fields in California, and preferably with grains or pulses as cash crops). She would have reached a very different conclusion using the global evidence (cover crops have neutral effects on cash crop yields). However, she found only three relevant studies, and there is some uncertainty in these results. It has been suggested that uncertainty could be incorporated into decision analysis (Gregory *et al.* 2012). She could use results of her dynamic meta-analysis — the mean effect size and its confidence interval — as inputs for decision analysis. However, we will leave this hypothetical user here, having shown some of the key features of dynamic meta-analysis on Metadataset.

## Discussion

Dynamic meta-analysis provides a partial solution to an important problem in evidence-based decision making — lack of access to relevant evidence (Christie *et al.* in press; Innvær *et al.* 2002; Cook, Possingham & Fuller 2013) — not only by helping users to find locally-relevant evidence in a global evidence base, but also by helping them to use this evidence to reach locally-relevant conclusions. We showed how the Metadataset website can be used for dynamic meta-analysis. For example, we showed how a hypothetical user could reach a different conclusion when using the global evidence (cover crops have no effect on cash crop yields) instead of the locally-relevant evidence (brassicas have negative effects on cash crop yields in California). As a next step, this evidence could be used as an input into decision analysis (Shackelford *et al.* 2019), but that is beyond the scope of our work here. Here we discuss some strengths and weaknesses of dynamic meta-analysis, and we suggest that this method could be scaled up and used for subject-wide evidence synthesis.

### Dynamic meta-analysis for subject-wide evidence synthesis

Metadataset was developed as part of the Conservation Evidence project (Sutherland *et al.* 2019), which provides summaries of scientific studies (including the studies of cover crops (Shackelford *et al.* 2017) that we used as an example of dynamic meta-analysis). By browsing and searching the Conservation Evidence website (www.conservationevidence.com), users may already be able to find summaries of studies that match their local conditions. In this sense, Metadataset does not represent progress beyond the interface that is already available on Conservation Evidence. However, Metadataset goes a step further. It enables users to reach new conclusions based on these studies.

This is only possible because Metadataset provides quantitative evidence (effect sizes) that can be dynamically reanalysed, whereas Conservation Evidence provides qualitative evidence (“effectiveness categories” (Sutherland *et al.* 2019)) that cannot yet be dynamically reanalysed. It is possible that dynamic methods could be developed for Conservation Evidence, perhaps by using expert assessment to assign quantitative scores to each study. However, there are good reasons that Conservation Evidence does not yet use quantitative methods. For example, the populations and outcomes of conservation studies are heterogeneous, and this suggests that meta-analysis might not be an appropriate method of evidence synthesis (Christie *et al.* in press), whereas agricultural studies may be more homogenous. Nevertheless, in subject areas for which quantitative methods are appropriate, Metadataset represents progress towards the co-assessment of evidence (Sutherland, Shackelford & Rose 2017), and dynamic meta-analysis complements the qualitative methods that are used by Conservation Evidence.

We suggest that dynamic meta-analysis could be particularly useful in the context of *subject-wide evidence synthesis* (Sutherland & Wordley 2018; Sutherland *et al.* 2019), which is a method of evidence synthesis that was developed by the Conservation Evidence project. Whereas a typical systematic review includes studies of only one or a few interventions, a subject-wide evidence synthesis includes studies of all interventions in a subject area (e.g., bird conservation), and thus it benefits from economies of scale (Sutherland & Wordley 2018). For example, a publication only needs to be read once, and all of the data can be extracted for all interventions, rather than needing to be read once for each review of each intervention.

Subject-wide evidence synthesis is evidence synthesis on the scale that is needed for multi-criteria decision analysis (Shackelford *et al.* 2019), and thus it is particularly relevant to a discussion of evidence-based decision making. Because subject-wide evidence synthesis is global in scale, it begs the question, “How relevant is this global evidence for my local decision?” We suggest that dynamic meta-analysis, or some similar method of assessing the local relevance of global evidence, could be especially useful for subject-wide evidence synthesis. On Metadataset, our work on invasive plant management (Martin *et al.* 2020) is an example of subject-wide evidence syntheses in conservation biology, and it will soon be possible to assess the transferability of this evidence using dynamic meta-analysis. It will also be possible to browse this evidence by intervention and outcome, and thus to consider its applicability to a specific decision (using dynamic meta-analysis only for those interventions and outcomes that are considered to be applicable).

### Metadataset compared to other tools

Researchers in psychology have suggested “community augmented meta-analysis” (CAMA), in which open-access databases of effect sizes could be updated and reused by researchers for future meta-analyses (Tsuji, Bergmann & Cristia 2014). MetaLab (http://metalab.stanford.edu) is an implementation of CAMA that includes data from several meta-analyses in psychology (Bergmann *et al.* 2018). It enables researchers to test the effects of covariates on the mean effect size (using meta-regression), but it does not provide options for subgroup analysis or recalibration. Metalab and other interactive databases of effect sizes could presumably be modified to provide these options. However, as we suggested above, dynamic meta-analysis could be particularly useful for subject-wide evidence synthesis, and therefore it would perhaps be better to have one large database for each subject, with interoperable data and metadata, rather than many small databases.

An older, offline tool that seems to be more similar to Metadataset in both function and intention is the Transparent Interactive Decision Interrogator (TIDI) in medicine (Bujkiewicz *et al.* 2011). TIDI provides options for subgroup analysis and study exclusion, but not recalibration. A newer, online tool is IU-MA (www.iu-ma.org), which provides “interactive up-to-date meta-analysis” of two datasets in medicine (Becker *et al.* 2020). Becker et al. (2020) also refer to dynamic meta-analyses, but they do not provide a definition of the term, and although their IU-MAs provide options for subgroup analysis, they do not provide options for recalibration.

All of these tools are clearly useful, and there are clearly many similarities between them, but there are also many differences. One important difference is that none of these tools, with the exception of Metadataset, provides options for recalibration (i.e. weighting individual studies based on their relevance) or for analysing the data at different levels of resolution (i.e., lumping or splitting interventions and outcomes before starting a dynamic meta-analysis). We see recalibration as a key feature for dynamic meta-analysis. We also see this lumping or splitting of evidence (which we will refer to as the *dynamic scoping* of a meta-analysis) as a key feature. As well as assessing the transferability of evidence using dynamic meta-analysis, we suggest that users should be able to assess the applicability of evidence by dynamically scoping the meta-analysis (which is also a process of filtering the evidence, like subgroup analysis, but it is done before starting the meta-analysis). Dynamic scoping could also provide a partial solution to the “apples and oranges” problem in meta-analysis (Sharpe 1997), since users could decide for themselves which “apples” and which “oranges” should be compared (e.g., deciding which interventions and/or outcomes should be analysed together). Therefore, we think that both filtering (subgroup analysis and dynamic scoping) and weighting (recalibration) should be seen as key features of dynamic meta-analysis. However, we note that both recalibration and dynamic scoping need to be further developed (see below).

Recalibration has the potential to improve evidence synthesis in subject areas where there is not any evidence that is completely relevant to decision makers (where subgroup analysis would not be useful). This relates to another important difference between these tools, which is that they are solutions to different problems, in different disciplines (agroecology, conservation biology, medicine, and psychology). In some disciplines, the need for recalibration may be less important than we perceive it to be in agroecology and conservation biology, in which there may be no evidence for a specific biome or taxon (Christie *et al.* in press; Cook, Possingham & Fuller 2013), and in which heterogeneity may be higher than it is in carefully controlled clinical or laboratory sciences. Thus, recalibration and other methods of assessing existing evidence may be especially important in disciplines with sparse evidence (cf. Sutherland & Wordley 2018).

### Protocols for evidence use

Dynamic meta-analysis could lead to a rebalancing of power and responsibility in evidence-synthesis, since evidence users would be able to make decisions that are typically made by researchers (Table 2). Protocols for evidence synthesis by researchers are well developed (e.g., CEE 2018), but protocols for evidence use by decision makers may need to be developed. Researchers who reanalyse existing datasets already need to take extra steps to avoid conflicts of interest and other perverse incentives (Christakis & Zimmerman 2013). However, these steps may become even more important as data is reanalysed not by researchers but by policy makers or other evidence users, especially if they have political agendas or other conflicts of interest that might bias their conclusions.

**Table 2.** Some comparisons between static and dynamic meta-analysis. In dynamic meta-analysis, many decisions are made by users, not researchers. However, these decisions are informed by researchers, who provide the metadata on which the decisions are based. In a static meta-analysis, most decisions are made by researchers. However, these decisions are often informed by users, who are often consulted when the protocol for a meta-analysis is being developed. Thus, both researchers and users can be involved in both static and dynamic meta-analysis, but only in dynamic meta-analysis can users interact with the methods and results.

| Questions | Static | Dynamic | Strengths (+) and weaknesses (-) of dynamic meta-analysis |
| --- | --- | --- | --- |
| Which interventions should be reviewed?<br>Which outcomes should be reviewed? | Researchers decide | Users decide | + Users can decide whether interventions and outcomes should be split or lumped (e.g., as comparisons of “apples and oranges”)<br>– Researchers may not have classified interventions and outcomes in a way that is relevant to users |
| Which studies should be included? High-quality studies only? Low-quality studies that are locally relevant? | Researchers decide | Users decide | + Users can include/exclude studies based on relevance and study quality<br>+ Users can weight studies based on relevance and study quality<br>– Users may not understand the limitations of study quality (e.g., blocking, controls, correlation vs causation, etc.)<br>– Researchers may not have classified study quality or described methods in a way that is relevant to users (poor reporting of methods or missing metadata) |
| Which results are informative? | Researchers decide | Users decide | + Users can explore results that researchers may not have explored (e.g., cover crops that are brassicas, in the USA, with irrigation)<br>– Users may not understand, or may be overwhelmed by, the analysis methods and results (e.g., multiple options)<br>– Researchers may not have classified metadata in a way that is relevant to users |
| Which results are credible? | Researchers decide | Users decide | + Users can select, deselect, and adjust settings to control the assumptions<br>+ Users can permute settings for sensitivity analysis<br>– Users may not understand the limitations of the analysis methods and results (e.g., model validity)<br>– Results may be vulnerable to cherry picking, data dredging, and other biases, if protocols for evidence use are not developed |

For example, if a user does multiple analyses, selecting and deselecting different filters, then it will be difficult to interpret the statistical significance of their results, because of the multiple hypothesis tests that this involves (the problem of “data dredging”) (Szucs 2016). Furthermore, if a user does multiple analyses, and selects only one of these analyses as the basis for their decision (perhaps because it supports their political agenda), then it will be difficult to defend the credibility of their conclusions (the problem of “cherry picking”). Protocols for evidence use could require dynamic meta-analyses to be predefined (e.g., predefining the filters that would be selected), and users could be restricted to a limited number of analyses.

### Standardized classification systems for metadata

Dynamic meta-analysis is limited by the quantity and quality of data and metadata that are available for each study. It has often been suggested that standards of data reporting need to be improved (e.g., Gurevitch & Hedges 1999), but here we suggest that standards of metadata reporting also need to be improved, and standardized systems for classifying metadata need to be developed for use in evidence synthesis. For Metadataset, we developed hierarchical classification systems for interventions and outcomes, and we will refine these systems as we review new studies. Standardized classification systems for other forms of metadata (e.g., terrestrial ecoregions (Olson *et al.* 2001)) will either need to be adopted or developed (e.g., as an extension of Ecological Metadata Language (Michener *et al.* 1997)). If a unified system could be developed for classifying all of the interventions, outcomes, and other metadata within a discipline, then the evidence from multiple subject-wide evidence syntheses could be integrated into a single discipline-wide database with interoperable data and metadata (cf. Sutherland *et al.* 2019). This should not be seen as a precondition for dynamic meta-analysis, but it could be a vision for the future.

## Conclusion

Nature is infinitely variable, and in many disciplines it is simply not possible to make generalizations that are universally applicable and transferable. But neither is it possible to be infinitely patient in waiting for locally-relevant evidence to be co-produced for every decision. If decisions need to be made quickly and efficiently, they may need to be based on the co-assessment of existing evidence, rather than the co-production of new evidence (Sutherland, Shackelford & Rose 2017). Here we have defined dynamic meta-analysis as a method that can be used for the co-assessment of existing evidence. We have also shown how this method could be used to reach new conclusions from existing evidence, with the example of Metadataset.

## Supporting information

Supplementary File 1 - Metadataset video

Supplementary File 3 - Examples of methods in R

Supplementary File 4 - Data for examples of methods in R

