## Supplementary File 2 - Supplementary methods for "Dynamic meta-analysis: a method of using global evidence for local decision making"

The Shiny app uses the methods from Shackelford et al. (2019) to calculate the log response ratio and its variance. We will not repeat these methods here, since Shackelford et al. (2019) is Open Access. However, we will review the assumptions that underpin these methods, and we will show how these assumptions can be changed, by selecting, deselecting, and/or adjusting the settings in the Shiny app.

Standard methods of calculating the variance of the log response ratio require standard errors and sample sizes, which are often unreported in research publications. It has been suggested that it is better to approximate or impute the missing data in a meta-analysis than it is to exclude the publications with missing data (Lajeunesse 2013). Sensitivity analysis can be used to test the assumptions that are used for approximation or imputation, as shown by Shackelford et al. (2019). If standard errors (or standard deviations) and sample sizes are unavailable, the Shiny app approximates the variance (*v*) of the log response ratio (*L*) using the *Z*-value (often calculated from the *P*-value), by using this equation.

|*L*| – (*Z* * *√v*) = 0

In other words, it uses the equation for the confidence interval, CI = *L* ± Z * *√v* (Hedges, Gurevitch & Curtis 1999), to set the lower or upper bound of the (1 – *P*) * 100% confidence interval to zero, and then it calculates *v* from this equation. In the dynamic meta-analysis of cover crops, this option enabled us to include many more publications. However, this option can be deselected using the checkbox for “Approximate the variance of the log response ratio using its (assumed) p-value or z-value”.

If exact *P*-values are unavailable, because they were reported as “significant” or “non-significant” (e.g., *P* < 0.05 or *P* > 0.05), then the Shiny app assumes an exact value of 0.025 for “significant” results and 0.525 for “non-significant” results. However, these default values can be adjusted via sliders in the Shiny app (and these values will not be used at all if the checkbox is deselected). If even *P*-values are unavailable, then variance can be imputed using the mean variance of all other included studies, but this can also be deselected using the checkbox for “Impute the variance for rows without variance (using the mean variance)”. If selected, the mean variance is calculating using a linear model with the same random effects as the meta-analysis, using the lme package in R.

The random effects in all models are specified as “random = ~ 1 | publication/study” (i.e. study is nested with publication). The “study” variable is dynamically generated by concatenating “study_ID” (which is statically defined by researchers in the Django app) and any other filter variables (which are dynamically select by users in the Shiny app). For example, Shackelford et al. (2019) defined “studies” as experiments with different species of cash crops and/or cover crops, even if these experiments were reported in the same publication. Thus, the user could select “Cash crop” and “Cover crop” in the “Random effects” section of the Shiny app. The Shiny app would then dynamically generate a new variable by pasting together “study_ID”, “Cash.crop” and “Cover.crop” (e.g., “Study ID 1 Maize Rye”) and then use this new variable as the “study” in the formula for random effects.

Studies with exceptionally high variance (*outliers*) can be defined in terms of deviations from the median variance (*median absolute deviance* or *MAD* (Leys et al. 2013)), and there is a slider for this in the Shiny app. Outliers can be excluded from the analysis, and there is a checkbox for this. The default setting is to exclude outliers, but the default threshold for defining outliers is relatively inclusive (10 deviations from the median variance). Excluding outliers can sometimes solve problems with convergence failures in the metafor model, which would otherwise show as error messages, and this relatively inclusive threshold for excluding outliers seems to be a useful default setting.

We think these default settings represent reasonable assumptions, but these settings can be selected, deselected, and/or adjusted, and sensitivity analysis can be used to test the effects of these assumptions. If users need more control than this, then they can download a CSV file and analyse the data themselves using R or other software packages.
