## Supplementary File 3 - Examples of methods in R for "Dynamic meta-analysis: a method of using global evidence for local decision making"

# This code was tested in R version 3.6.3 with metafor version 2.4-0 and MuMIn version 1.43.17.
library(metafor)
library(MuMIn)
eval(metafor:::.MuMIn)


###############################
# (1) Meta-analysis of all data
###############################

# This script uses the CSV file on crop yield that was published as supplementary material. Download 
# the CSV file and load it into R (316 data points on cash crop yield). To follow this example, change 
# "path/to/filename.csv" to that of the downloaded CSV file on your computer.
data <- read.csv("/path/to/filename.csv")
data <- data[c("citation", "publication", "study", "log_response_ratio", "selected_v", "Country", 
  "Cover.crop.type", "Irrigated.cash.crop", "relevance_weight")]
dim(data)   # 316 rows of data
head(data)  # The first 6 rows of data

# Meta-analysis of all data (without filters). In the Shiny app, this is the data with the checkbox 
# deselected for "Exclude rows with exceptionally high variance (outliers)" (i.e. this data includes 
# outliers).
meta_analysis <- rma.mv(
  log_response_ratio,
  selected_v,
  random = ~ 1 | publication/study,
  data = data
)
summary(meta_analysis)

# The results are back-transformed from log response ratio to response ratio and rounded.
round(exp(meta_analysis$b), 2)      # Response ratio
round(exp(meta_analysis$ci.lb), 2)  # Lower bound of the 95% confidence interval
round(exp(meta_analysis$ci.ub), 2)  # Upper bound of the 95% confidence interval
round(meta_analysis$pval, 4)        # P-value


#######################
# (2) Subgroup analysis
#######################

# Subset the data.
subgroup_data <- subset(data, 
  Country == "United States of America" &
  Cover.crop.type == "Brassica" &
  Irrigated.cash.crop == "Yes"
)
dim(subgroup_data)  # 14 rows of data from 2 publications

subgroup_analysis <- rma.mv(
  log_response_ratio,
  selected_v,
  random = ~ 1 | publication/study,
  data = subgroup_data
)
summary(subgroup_analysis)

# The results are back-transformed from log response ratio to response ratio and rounded.
round(exp(subgroup_analysis$b), 2)      # Response ratio
round(exp(subgroup_analysis$ci.lb), 2)  # Lower bound of the 95% confidence interval
round(exp(subgroup_analysis$ci.ub), 2)  # Upper bound of the 95% confidence interval
round(subgroup_analysis$pval, 4)        # P-value


#####################
# (3) Meta-regression
#####################

# Fit a full model with all variables and all two-way interactions for use in model selection.
full_model <- rma.mv(
  log_response_ratio, 
  selected_v,
  mods = ~ Country + Cover.crop.type + Irrigated.cash.crop 
    + Cover.crop.type:Irrigated.cash.crop + Country:Irrigated.cash.crop + Country:Cover.crop.type,
  random = ~ 1 | publication/study, 
  data = data,
  method = "ML"  # ML is needed for log-likelihood comparisons with MuMIn, but we will refit with REML.
)

# Use MuMIn for model selection.
model_selection <- dredge(full_model, trace = 2)
subset(model_selection, delta <= 2, recalc.weights = FALSE)

# Get the model with the lowest AICc and refit with REML.
best_model <- get.models(model_selection, subset = 1, method = "REML")[[1]]
summary(best_model)

# Get the moderators that were included in the best model. "Irrigated.cash.crop" and all interactions
# were excluded from the best model in this example, but "Country" and "Cover.crop.type" were included. 
best_model$formula.mods

# Get the model predictions for brassicas in the USA.
meta_regression <- predict(best_model, newmods = c(0,0,0,1,1,0,0,0))
meta_regression

# The results are back-transformed from log response ratio to response ratio and rounded.
round(exp(meta_regression$pred), 2)                     # Response ratio
round(exp(meta_regression$ci.lb), 2)                    # Lower bound of the 95% confidence interval
round(exp(meta_regression$ci.ub), 2)                    # Upper bound of the 95% confidence interval
zval <- abs(meta_regression$pred) / meta_regression$se  # Z-value
pval <- 2 * (1 - pnorm(zval))
pval <- round(pval, 4)
pval                                                    # P-value

# Because "Irrigated.cover.crop" was not included in the best model, the user decides to remove this
# filter and redo the subgroup analysis.
subgroup_data <- subset(data, 
  Country == "United States of America" &
    Cover.crop.type == "Brassica"
)
dim(subgroup_data)  # 30 rows of data from 3 publications

subgroup_analysis <- rma.mv(
  log_response_ratio,
  selected_v,
  random = ~ 1 | publication/study,
  data = subgroup_data
)
summary(subgroup_analysis)


###################
# (4) Recalibration
###################

# M is the marginal variance-covariance matrix.
M <- subgroup_analysis$M
M

# W is weight matrix (inverse-variance weights), which is the inverse of M.
W <- solve(M)
W

# We get the same results if we explicitly specify this weight matrix with W = W.
subgroup_analysis_W <- rma.mv(
  log_response_ratio, 
  selected_v, 
  W = W, 
  random = ~ 1 | publication/study,
  data = subgroup_data
)
summary(subgroup_analysis)
summary(subgroup_analysis_W)

# In this example, the user assigns a relevance weight of 0.5 to Hartz et al., 2005.
citations <- unique(subgroup_data$citation)
Hartz_et_al_2005 <- citations[2]
Hartz_et_al_2005
subgroup_data$relevance_weight[subgroup_data$citation == Hartz_et_al_2005] <- 0.5

# C is a diagonal matrix based on the relevance weights.
C <- diag(subgroup_data$relevance_weight)
C

# As a provisional example of recalibration, we modify the weight matrix, W, using the matrix of 
# relevance weights, C.
WC <- sqrt(C) %*% solve(M) %*% sqrt(C)
WC

# We specify W = WC in the subgroup analysis.
subgroup_analysis_WC <- rma.mv(
  log_response_ratio, 
  selected_v, 
  W = WC, 
  random = ~ 1 | publication/study, 
  data = subgroup_data
)
summary(subgroup_analysis_WC)

# This is the proportional weight assigned to each study in the recalibrated subgroup analysis.
weights <- solve(t(subgroup_analysis_WC$X) %*% subgroup_analysis_WC$W %*% subgroup_analysis_WC$X) %*% 
  t(subgroup_analysis_WC$X) %*% subgroup_analysis_WC$W
weights

# These proportional weights can be checked against the model results.
# Log response ratio:
subgroup_analysis_WC$b
weights %*% subgroup_analysis_WC$yi
# Standard error of the log response ratio:
subgroup_analysis_WC$se
sqrt(weights %*% subgroup_analysis_WC$M %*% t(weights))

# The results are back-transformed from log response ratio to response ratio and rounded.
round(exp(subgroup_analysis_WC$b), 2)      # Response ratio
round(exp(subgroup_analysis_WC$ci.lb), 2)  # Lower bound of the 95% confidence interval
round(exp(subgroup_analysis_WC$ci.ub), 2)  # Upper bound of the 95% confidence interval
round(subgroup_analysis_WC$pval, 4)        # P-value
